# Associations between lower limb isometric torque, isokinetic torque, and explosive force with phases of reactive stepping in young, healthy adults

**DOI:** 10.1101/2021.01.18.427144

**Authors:** Tyler M. Saumur, Jacqueline Nestico, George Mochizuki, Stephen D. Perry, Avril Mansfield, Sunita Mathur

**Author notes:** **Correspondence:** Tyler Saumur, MSc, PhD candidate, Rehabilitation Sciences Institute, University of Toronto, 160-500 University Ave, Toronto ON M5G 1V7, Canada.

## Abstract

Reactive stepping is one of the only strategies that can lead to successful stabilization following a large challenge to balance. Improving function of specific muscles associated with reactive stepping may improve features of reactive balance control. Accordingly, this study aimed to determine the relationship between lower limb muscle strength and explosive force with force plate-derived timing measures of reactive stepping. Nineteen young, healthy adults (27.6 ± 3.0 years of age; 10 women: 9 men) responded to 6 perturbations (~13-15% of body weight) using an anterior lean-and-release system (causing a forward fall), where they were instructed to recover balance in as few steps as possible. Foot-off, swing, and restabilization times were estimated from force plates. Peak isokinetic torque, isometric torque, and explosive force of the knee extensors/flexors and plantar/dorsiflexors were measured using isokinetic dynamometry. Correlations were run based on *a priori* hypotheses and corrected for the number of comparisons (Bonferroni) for each variable. Knee extensor explosive force was negatively correlated with swing time (r = −0.582, p = 0.009). Knee flexor peak isometric torque also showed a negative association with restabilization time (r = −0.459, p = 0.048), however this was not statistically significant after correcting for multiple comparisons. There was no significant relationship between foot-off time and knee or plantar flexor explosive force (p > 0.025). These findings suggest that there may be utility to identifying specific aspects of reactive step timing when studying the relationship between muscle strength and reactive balance control. Exercise training aimed at improving falls risk should consider targeting specific aspects of muscle strength depending on specific deficits in reactive stepping.

## 1. Introduction

Reactive balance control is necessary for maintaining or regaining stability following a balance perturbation (Shumway-Cook & Woollacott, 2012). Reactive balance control involves the activation of trunk and lower limb muscles (Chvatal et al., 2011; Chvatal & Ting, 2013). Accordingly, muscle strength is commonly studied in the context of balance control. Muscle strength can be defined as the force-generating capacity of the muscle (Robles et al., 2011) and can be quantified using various measures of force or torque. Muscle power is the mechanical work performed by the muscle over time and the muscle’s ability to produce high velocity movements (Sapega & Drillings, 1983). Balance control, strength, and power decline as we age (Baloh et al., 2003; Ferretti et al., 1994; Hughes et al., 2001); however, variability in balance, strength, and power levels is even observed in the young adult population (Laforest et al., 1990; Petroman & Rata, 2020; Zemková et al., 2016, 2017, 2019). It might be expected that strength and reactive balance control would be related to one another as our muscles play an important role in recovering balance. However, there is limited evidence supporting this relationship in young adults.

In a systematic review exploring the relationship between strength, power and balance control (Muehlbauer et al., 2015), only three studies incorporated reactive/dynamic balance components when studying young adults (Izquierdo et al., 1999; Muehlbauer et al., 2012; Piirainen et al., 2010). In these studies, muscle power was modestly correlated (weighted mean *r*_z_ of 0.27, *p* = 0.006) with reactive balance performance, whereas explosive force did not show a significant correlation (weighted mean *r*_z_ of 0.26, *p* = 0.09). Similarly, a recent study found that neither explosive force nor peak isometric torque was associated with centre of pressure (COP) variables during foot-in-place perturbations in sedentary and physically active young adults (Zemková et al., 2017). In these studies, perturbations were not large enough to elicit a reactive step; increasing perturbation magnitude results in larger muscular responses and higher joint torques (Thelen et al., 2000; Wojcik et al., 2001). Thus, larger sized perturbations may be required to expose potential associations between strength and balance control. Following lean-and-release perturbations, which evoke a stepping response, Wojcik and colleagues (2001) found that peak isometric torque of the plantar flexors and hip flexors were poor predictors of reactive stepping performance. Conversely, Karamanidis et al. (2008) found relationships between peak isometric plantar flexor and knee extensor torque and margin of stability following a forward loss of balance. More recently, explosive force was found to be associated with reactive balance performance in young adults (Ochi et al., 2020). In particular, hip and knee flexor explosive force explained 59% of the variance in reactive balance performance.

The variable findings in studies implementing reactive stepping paradigms may in part be due to the protocol and measures used for assessing reactive balance performance. Wojcik et al. (2001) and Ochi et al. (2020) both used maximum lean angle as their primary measure of reactive balance control. Maximum lean angle is a measure of the largest perturbation magnitude at which individuals can successfully regain stability in a single step following a lean-and-release perturbation. Conversely, Karamanidis and colleagues (2008) used margin of stability – a measure which reflects the capacity of the lower limbs to maintain postural stability. While these measures have been effective in revealing relationships between muscle strength and reactive stepping, there may be an added benefit to investigating reactive step timing in relation to muscle strength as the functional roles of lower limb muscles differ throughout the reactive stepping response. Specific aspects of reactive stepping such as foot-off time, swing time, and time to restabilization have been commonly reported and are related to response success and falls (Brauer et al., 2001; Maki & McIlroy, 1997; Mansfield et al., 2015). It is likely that aspects of muscle strength and power are related to the duration of reactive balance phases; however, to our knowledge, work to date has not investigated the relationship between muscle strength and reactive stepping phases.

The purpose of this study was to determine the relationship between lower limb muscle strength and explosive force (a surrogate of muscle power) with force plate-derived timing measures of reactive stepping. Based on previous electromyography data (Thelen et al., 2000) and previous muscle strength studies (Karamanidis et al., 2008; Ochi et al., 2020), hypotheses were generated taking into consideration the function of the muscles and the amplitude and slope of their muscle activity during the reactive stepping phases. Accordingly, it was hypothesized that (1) knee flexor and ankle plantar flexor explosive force would be negatively associated with foot-off time; (2) knee extensor explosive force and isokinetic peak torque would be negatively associated with swing time; and (3) knee flexor and ankle plantar flexor peak isometric torque would be associated with time to restabilization.

## 2. Methods

### 2.1. Participants

Participants who met the following inclusion criteria were enrolled in the study: between the ages of 20 and 35 years old, able to comprehend English, no self-reported walking difficulties, no neurological or musculoskeletal disorders, normal or corrected to normal vision, and no conditions that limit one’s ability to complete daily activities. Participants were recruited from the Greater Toronto Area and research ethics approval was obtained from Sunnybrook Health Sciences Centre (#017-2015) and the University of Toronto (#31805). All participants provided written, informed consent. Individuals who participated in this study were also part of another study on reliability of reactive stepping measures. Balance data from the second data collection session was used and has been partially reported elsewhere (Saumur et al., 2021).

### 2.2. Muscle Strength Assessment

Strength testing of the knee extensors/flexors and ankle plantar/dorsiflexors was performed using a HUMAC NORM dynamometer (CSMi Medical Solutions, Stoughton, MA) on the preferred stepping limb determined by the reactive balance control task which corresponded to the dominant support foot in 15 of 19 participants, as determined by the Waterloo Footedness Questionnaire (Elias et al., 1998). For all strength testing, participants were instructed to “push/pull as hard and fast as possible” and were verbally encouraged throughout each repetition. No visual feedback of torque output was provided. The testing limb was firmly strapped to the dynamometer arm. Hip and shoulder straps were used to secure and position individuals in the chair, as appropriate. Mechanical stops were put in place to ensure participants’ safety. Isokinetic testing of the limbs was performed first, followed by isometric muscle testing. A rest period of 1 minute was provided in between each set and test.

#### 2.2.1. Isokinetic Testing

Isokinetic testing at both joints was performed at 180°/s. Isokinetic muscle strength for the knee and ankle at this velocity has been previously explored in the context of balance control in various populations (Lanzarin et al., 2016; Lehnert et al., 2017) and also mimics joint velocities during reactive stepping (Luchies et al., 1994). For isokinetic testing, participants initially performed a warm-up set of five repetitions at approximately 50% maximal effort.

Following a 1-minute rest period, participants performed a maximal effort single test set of 5 repetitions. For knee extension and flexion, participants were in a seated position at a hip angle of 85° with their hands across their chest. Participants started with their knee in a flexed position and performed 5 repetitions of knee extension and flexion in succession. For isokinetic ankle torque, participants were seated with a hip angle of 85° and a knee angle of 180° (full extension) with their arms folded across their chest. Participants started with their foot in a dorsiflexed position and performed 5 repetitions of ankle plantar- and dorsiflexion in succession.

#### 2.2.2. Isometric Testing

For isometric strength testing, all joint angles for isometric muscle testing corresponded with muscle lengths that result in maximal muscle activity and torque generation (Arampatzis et al., 2006; Dalton et al., 2013; Kulig et al., 1984; Onishi et al., 2002; Sale et al., 1982). Participants were instructed to “push (or pull) as hard as fast as possible.” These instructions were given to enable the measurement of the rate of torque development as well as peak isometric torque. Participants performed an initial warm up trial at approximately 50% maximal effort which was held for 5 seconds. Participants then performed 3-5 MVICs that were held for 5 seconds, with 1-minute rest periods between each repetition. The number of repetitions varied depending on the number of repetitions needed to obtain 3 maximum values within 10% of each other. Knee extension was tested in a seated position (hip angle of 85°) with the knee at 120° (60° flexion) and arms crossed on the participant’s chest. For knee flexion, participants were placed in a prone position with a knee angle of 150° (30° flexion) and a 180° hip angle. For plantar flexor and dorsiflexor isometric testing, participants were seated similar to the isokinetic testing (hip angle of 85°, knee angle of 180°, arms across chest). For plantarflexion trials, the foot was positioned at 5° dorsiflexion (0° equivalent to a neutral ankle angle). For dorsiflexion trials, the foot was positioned at 10° plantarflexion.

### 2.3. Reactive Balance Control Assessment

To study reactive balance control, lean-and-release testing was performed (Inness et al., 2015; Figure 1). Participants stood on a single force plate (Type 9260AA, Kistler, Winterthur, Switzerland) with heel centres 17 cm apart and externally rotated 14° (McIlroy & Maki, 1997). Participants were harnessed with a cable to an in-line load cell (Transducer Techniques, Temecula, USA) behind them. Perturbations were evoked by randomly releasing the rear cable while participants were leaning forward, resulting in a stepping response. Participants leaned forward at the ankle joint with ~8-10% of their body weight supported by the posterior tether cable for the small perturbation or ~13-15% of body weight support for the large perturbation.

**Figure 1.**
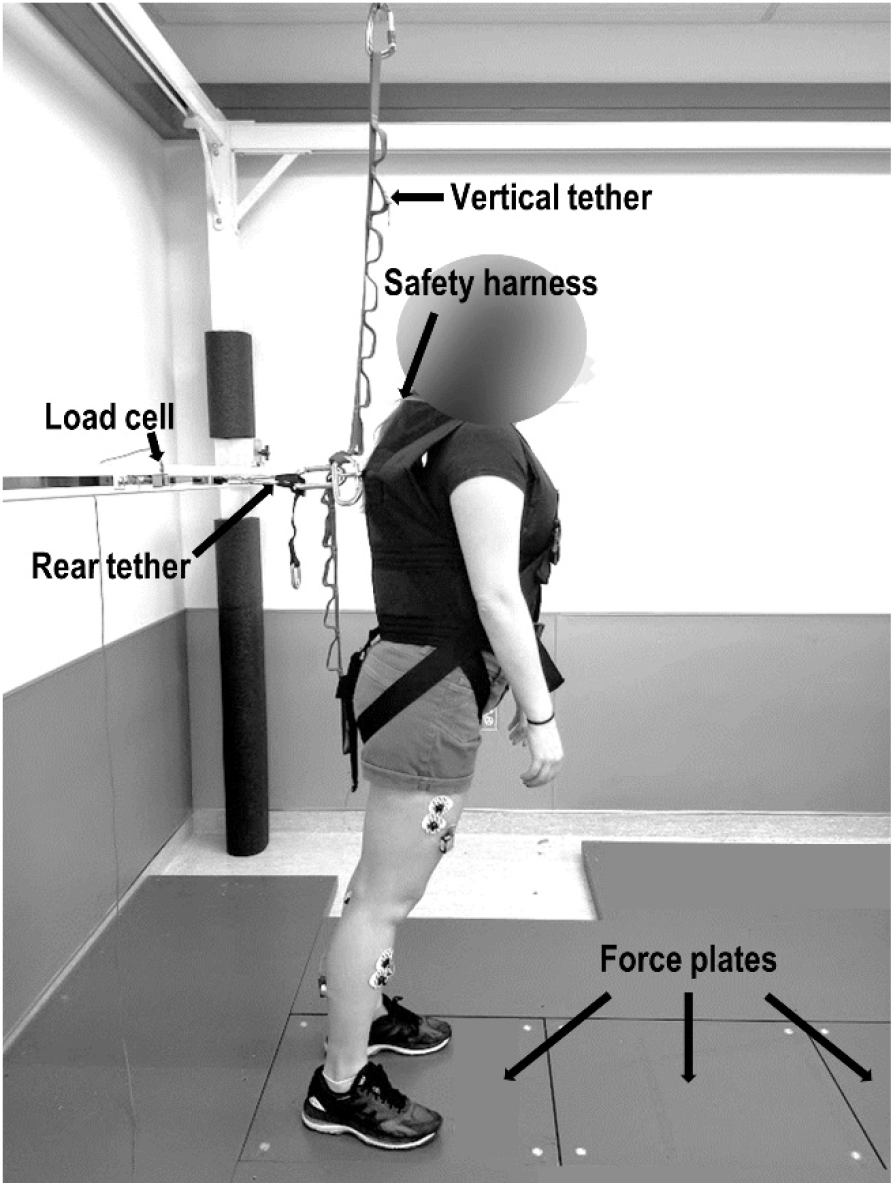
Example of experimental setup.

Participants were told to “recover their balance in as few steps as possible” with a pre-specified leg. Participants performed one small perturbation trial at the beginning of the session with no instructions regarding which foot to step with; this trial was excluded from analysis and the foot used was referred to as the “preferred stepping limb”. Participants performed a block of six trials for each leg. Perturbation size was randomized for each trial block, with each block consisting of three small and three large perturbations (12 total). After each perturbation, participants were instructed to maintain their position for 7 s to provide sufficient time to identify the stabilization of the COP velocity. Only the large perturbation condition data were used for analysis.

### 2.4. Questionnaires

The Activities-Specific Balance Confidence (ABC) Scale (Powell & Myers, 1995), International Physical Activity Questionnaire (IPAQ) short form (Craig et al., 2003), and Waterloo Footedness Questionnaire (Elias et al., 1998) were completed to determine the participants’ balance confidence, physical activity levels, and lower limb dominance, respectively. For the Waterloo Footedness Questionnaire, the balance task questions (question numbers 2, 4, 6, 8, and 10) were used for classifying the dominant support limb.

### 2.5. Data Analysis

Torque and joint angles were recorded at 100 Hz. Peak torque was obtained from the average of the 3 values within 10% of each other for each strength test. Explosive force was measured from the isometric trials as the rate of torque development from the time between 40-80% of peak torque (Webber & Porter, 2010). The trial with the highest calculated rate of torque development was used for analysis. Torque values were gravity-corrected and peak torques were obtained via custom code in Spike2 (Version 7.17, Cambridge Electronic Design, Cambridge, UK) and visually confirmed. Ground reaction forces and load cell data were sampled at 256 Hz and stored for offline analysis. Data were collected using a Power 1401 mkII (Cambridge Electronic Design, Cambridge, UK) and Spike2 data acquisition software. A schematic of the force plate-derived measures is in Figure 2. Foot-off time was from perturbation onset (< 5 N on load cell) to when the mediolateral COP was 20% of the max slope of the COP signal towards the stepping limb (Kurz et al., 2013). Swing time occurred from foot-off to foot contact (>5 N on stepping plate). Time to restabilization occurred from foot contact to the time at which the net anteroposterior COP velocity entered within 3 standard deviations of the mean net anteroposterior COP velocity – calculated from the last 2 s of foot contact – and remained for at least 1 s.

**Figure 2A.**
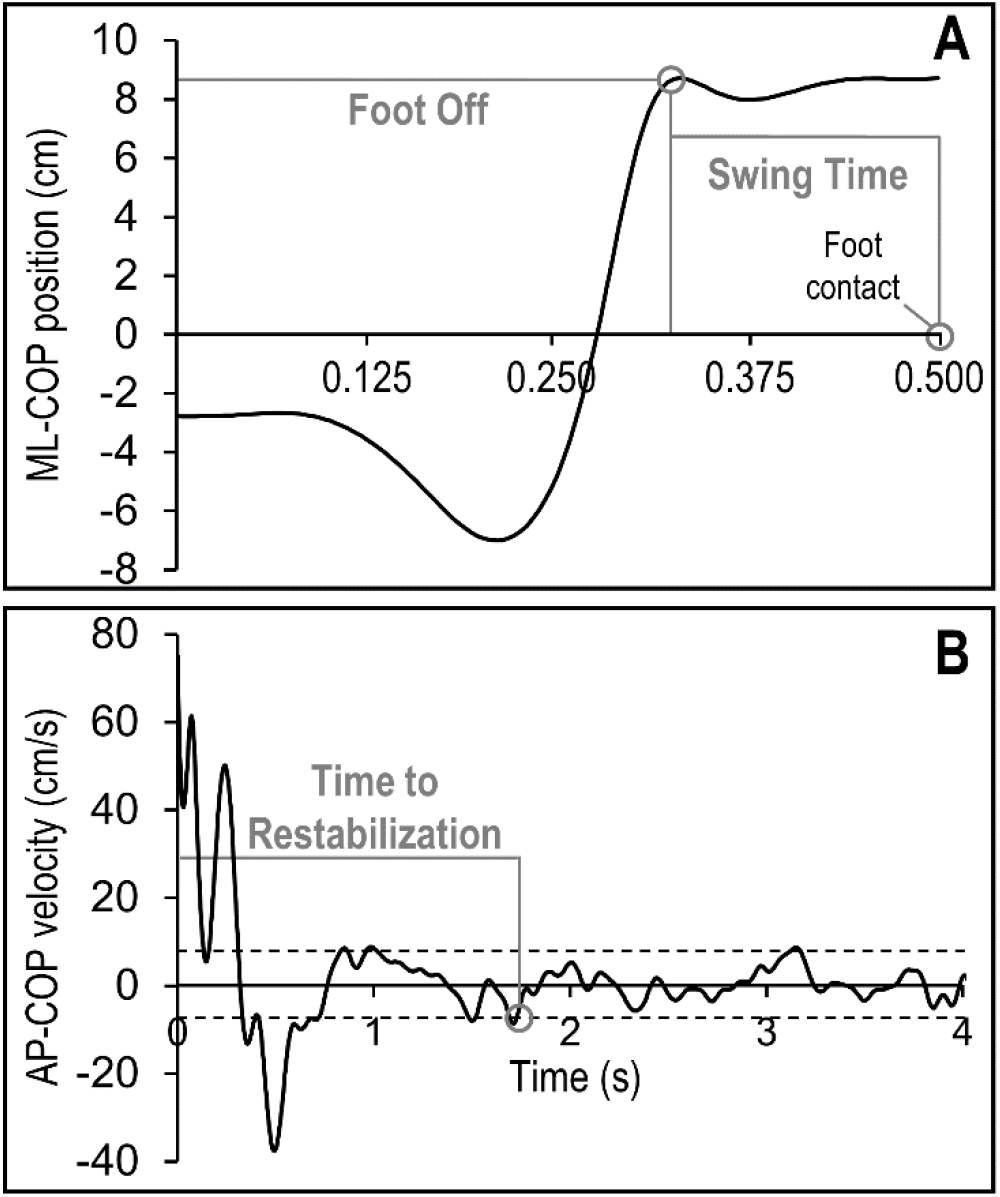
ML-COP position from perturbation onset to foot contact. **B.** Net AP-COP velocity Time starts at foot contact. ML-COP, mediolateral centre of pressure; AP-COP, anteroposterior centre of pressure.

### 2.6. Statistical Analysis

Statistical analysis was performed using SPSS Statistics version 26 (IBM, Armonk USA). Values are reported as mean (standard deviation) unless otherwise stated. Pearson correlations were performed for all hypothesized relationships between the force plate and muscle strength variables. As previous work has suggested that stronger relationships are present between muscle strength and reactive balance during larger perturbation magnitudes (Karamanidis et al., 2008), only the large perturbation force plate measures were used for analysis. This was also done to minimize the chance of a Type I error. To control for multiple correlations with each variable, a Bonferroni correction was performed, and significance was set to *p* < 0.025.

**Table 1.** Mean and standard deviations of measures from isokinetic and isometric testing

|  | Knee extensor | Knee flexor | Plantar flexor | Dorsiflexor |
| --- | --- | --- | --- | --- |
| Peak isokinetic torque | 96.3 (31.2) | 56.6 (19.9) | 43.0 (12.1) | 22.4 (8.7) |
| Peak isometric torque | 149.6 (47.0) | 43.8 (16.0) | 101.8 (28.4) | 36.4 (10.1) |
| Explosive force | 377.1 (237.5) | 98.7 (64.2) | 223.3 (106.7) | 123.4 (39.2) |

Units for peak isokinetic and isometric torque = Nm; units for explosive force = Nm/s

## 3. Results

Nineteen participants took part in this study (27.6 (3.0) years; 10 women and 9 men) as previously described (Saumur et al., 2021). Mean foot-off, swing, and restabilization time were 306.1 s (25.0 s), 168.1 s (27.3 s), and 1906.4 s (489.0 s), respectively. There were negative correlations between strength and timing measures (Figure 3), with higher strength associated with faster step times. However, for Hypothesis 1, neither knee flexor (*r* = −0.407, *p* = 0.084) nor ankle plantar flexor explosive force (*r* = −0.323, *p* = 0.178) were significantly correlated with foot-off time. Regarding Hypothesis 2, knee extensor explosive force (*r* = −0.582, *p* = 0.009), but not knee extensor isokinetic peak torque (*r* = −0.259, *p* = 0.284), was negatively correlated with swing time. For Hypothesis 3, there were no significant correlations between time to restabilization and knee flexor (*r* = −0.459, *p* = 0.048) or ankle plantar flexor peak isometric torque (*r* = −0.425, *p* = 0.070) after controlling for multiple comparisons.

**Figure 3.**
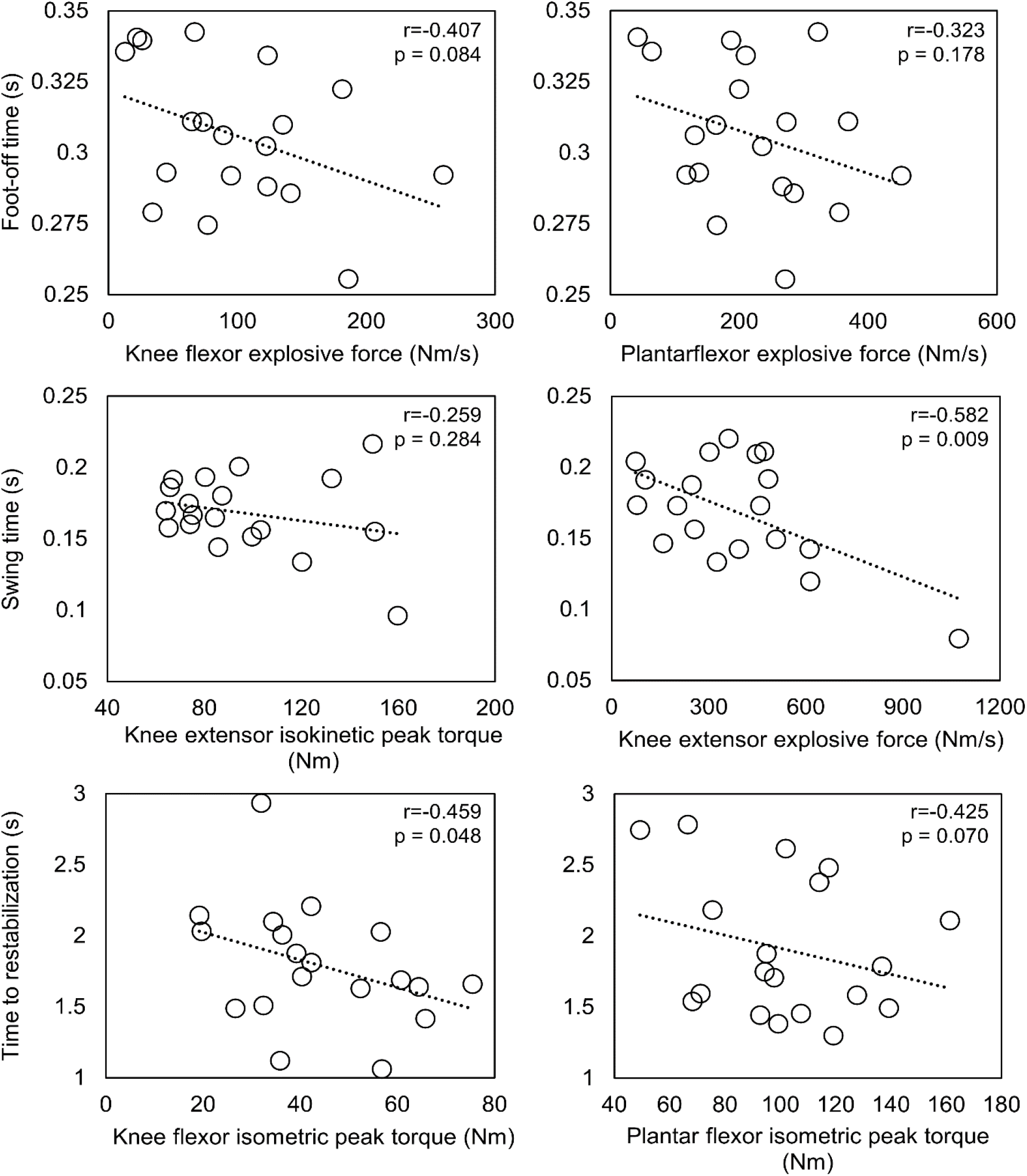
Scatter plots of the correlations performed between muscle strength and reactive step timing measures.

## 4. Discussion

This study found that all relationships between reactive step timing and muscle strength trended in the direction supporting increased strength being associated with faster phases of reactive stepping. Of note, knee extensor explosive force was moderately correlated to swing time. Knee flexor isometric strength – while moderately correlated with time to restabilization – was not statistically significant after adjusting for multiple comparisons (*r* = −0.459, *p* = 0.048). These findings suggest that there may be utility to identifying specific aspects of reactive step timing such as foot-off, swing, and restabilization time when studying the relationship between muscle strength and reactive balance control.

A statistically significant relationship between knee extensor explosive force and swing time was found in the present study. This contrasts previous work that showed knee extensor explosive force was not correlated with maximum lean angle during the lean-and-release test (Ochi et al., 2020). It may be that the temporal or spatial components of reactive stepping are more sensitive to detecting relationships between strength and balance control than a global measure of stability. Indeed, knee extensor isometric torque has been shown to be associated with margin of stability (Karamanidis et al., 2008). However, another potential explanation may be that the relationship of knee extensor explosive force with reactive balance performance depends on perturbation size. In our study, the large perturbation trials used for analysis corresponded to only 13-15% of participants body weight, which equates to a lean angle of approximately 13.5° (Thelen et al., 1997). Conversely, the maximum recoverable lean angle for participants in the study by Ochi et al (2020) was 32.4±5°. As swing time does not appear to change greatly with increased lean angle after ~15° (Hsiao & Robinovitch, 1999), knee extensor explosive force may play less of a role in stabilizing at higher lean angles which may explain why no association was observed with perturbations performed at maximum lean angle previously.

No relationship between isokinetic knee extensor strength and swing time was observed in the present study. We tested isokinetic knee extensor strength at 180°/s, as it was previously shown to be associated with static balance control in middle-aged and older adults (Lanzarin et al., 2016; Lehnert et al., 2017). In these studies, isokinetic knee extensor peak torque was correlated with mean anteroposterior COP velocity in quiet standing (Lehnert et al., 2017) and was significantly different between groups that did and did not have reactive balance control deficits (Lanzarin et al., 2016). Furthermore, mean changes in knee joint angle over the swing phase (Luchies et al., 1994) correspond with a joint velocity of 180°/s. However, while isokinetic strength testing is performed at a constant velocity, joint accelerations are variable throughout the swing phase of reactive stepping (Graham et al., 2014). Thus, there may be limited transfer of isokinetic strength testing to reactive step timing performance.

We initially hypothesized that knee flexor and ankle plantar flexor explosive force would be correlated with foot-off time. This hypothesis was based on the functional role of the knee flexors and ankle plantar flexors of the stepping limb for propelling the centre of mass forward through ankle flexion and hip extension (Graham et al., 2017). In addition, the steep rise in muscle activity of these muscle groups (Thelen et al., 2000) suggests a temporal urgency during activation (Corcos et al., 1989; Maffiuletti et al., 2016). Our findings differ from those of Ochi et al. (2020), who found a significant relationship between knee flexor explosive force and maximum lean angle. An explanation as to why no relationship was found with foot-off time may be that foot-off time is associated with voluntary reaction time, which was not measured in our study. Indeed, Lord & Fitzpatrick, (2001) found that both lower limb strength and simple reaction time were associated with choice stepping reaction time performance. Thus, while muscle strength may play a role in foot-off time, processing sensory information and relaying commands to the muscle effectors likely also plays an important role.

Lastly, while there were moderate correlations between time to restabilization and plantar flexor and knee flexor peak isometric torque, these relationships were not statistically significant. Peak isometric torque was selected for correlation analysis as peak isometric plantar flexor torque (Karamanidis et al., 2008) has been found to be associated with margin of stability following a perturbation, and both plantar flexors and knee flexors have increased muscle activation at foot contact with larger perturbation magnitudes (Saumur et al., 2020). As there is minimal change in knee and ankle joint velocity shortly after foot contact (Madigan & Lloyd, 2005), and stepping limb muscle activity following foot contact is not as steep as those observed at foot-off (Thelen et al., 2000), peak isometric torque was assessed. While peak isometric plantar flexor torque was previously found to be associated with margin of stability (Karamanidis et al., 2008), it was also found to not be associated with maximum lean angle (Wojcik et al., 2001). To our knowledge, peak knee flexor isometric torque has not been explored in the context of reactive balance control. Ochi et al. (2020) did however find a significant correlation between knee flexor explosive force and maximum lean angle, however this measure does not contain any temporal features; thus, the functional role of knee flexor explosive force on reactive stepping is uncertain. Another measure of muscle strength that may be associated with restabilization time is knee flexor eccentric strength of the knee flexors. Eccentric muscle activity and moments have been observed at foot contact (Nagano et al., 2015; Wu et al., 2007) and with maximum knee flexion occurring after foot contact (Nagano et al., 2015), eccentric knee flexor strength may be related to restabilization time. While aspects of knee flexor and ankle plantar flexor strength may be related to time to restabilization, we were unable to confirm this relationship within the present study.

### 4.1. Limitations

We failed to find statistically significant correlations in many of our analyses, with *p-* values approaching 0.05. The sample size of this study was comparable to others which both did, (Karamanidis et al., 2008) and did not (Piirainen et al., 2010; Wojcik et al., 2001), observe relationships between muscle strength and reactive balance performance in young adults. However, in order to have a sufficient sample size to observe statistically significant effects using a hypothesized r = −0.27 (Muehlbauer et al., 2015), a sample of approximately 125 individuals would be needed when controlling for multiple comparisons with 80% power. It would not have been feasible for us to recruit this many participants with our current resources. A second limitation to our study is that MVICs were performed in a specific seated or prone position at a pre-specified angle based on optimal muscle length and joint angle for peak torque production. These static positions do not necessarily reflect the functional angles of the muscles during reactive stepping and therefore may limit the transferability of the strength values to reactive stepping control. Future work may want to explore how force generation at the specific joint angles of foot-off or foot contact relate to performance in these variables. Lastly, eccentric muscle work has been observed at foot contact (Nagano et al., 2015; Wu et al., 2007) and eccentric muscle strength, particularly in the knee flexors, may therefore be related to time to restabilization. Future work should explore how eccentric muscle strength relates to restabilization time and other features of stability following foot contact.

### 4.2. Conclusions

This study explored the relationship between lower limb muscle strength and explosive force with force plate-derived timing measures of reactive stepping. We found that knee extensor explosive force was related to swing time. While there may be a benefit to identifying specific aspects of reactive step timing when studying the relationship between muscle strength, power and reactive balance control, future work should explore these relationships in larger sample sizes and in populations with balance deficits.

## Acknowledgements

The authors would like to thank all participants for their time and commitment.

